# Computational analysis reveals similarities and differences between SCLC subtypes

**DOI:** 10.1101/2021.10.27.465593

**Authors:** Abhay Singh, Parth Desai, Maalavika Pillai, Nilay Agarwal, Nobuyuki Takahashi, Anish Thomas, Mohit Kumar Jolly

**Affiliations:** Centre for BioSystems Science and Engineering, Indian Institute of Science, Bangalore, India; Department of Biosciences and Bioengineering, Indian Institute of Technology Guwahati, Guwahati, India; Developmental Therapeutics Branch, Center for Cancer Research, NCI, NIH, Bethesda, USA; Department of Medical Oncology, National Center for Global Health and Medicine, Tokyo, Japan

**Keywords:** SCLC, Neuroendocrine, Phenotypic heterogeneity, ASCL1, NEUROD1, POU2F3, YAP1, EMT

## Abstract

Small cell lung cancer (SCLC) is a neuroendocrine malignancy with dismal survival rates. Previous studies have revealed inter and intra tumoral heterogeneity of SCLC driven by neuroendocrine differentiation and multiple gene expression signatures have been proposed to classify the distinct SCLC molecular subtypes However, few questions remain unanswered: a) how many SCLC subtypes exist? b) how similar or different are these subtypes?, c) which gene list(s) can be used to identify those specific subtypes? Here, we show that irrespective of the three gene sets (33 genes, 50 genes, 105 genes) proposed in different studies to classify SCLC into different subtypes, the markers of phenotypic heterogeneity in SCLC form a “teams” like pattern of co-expressed modules. Moreover, the 105 geneset could classify SCLC cell lines into five clusters, three of which can be distinctly mapped to the SCLC-A, SCLC-N and SCLC-Y subtypes. Intriguingly, we noticed a high degree of similarity in the transcriptional landscape of two non-neuroendocrine subtypes: SCLC-Y and SCLC-I*, as well as in their enrichment of EMT. Thus, our analysis elucidates the landscape of phenotypic heterogeneity enabling diverse SCLC subtypes.

## Introduction

Small cell lung cancer (SCLC) is a highly aggressive neuroendocrine tumor of airway epithelium with an abysmal five-year survival rate of 7% (Gazdar *et al*., 2017). It accounts for roughly 15% of all lung malignancies, is strongly associated with smoking, has a proclivity for early metastasis, and a poor prognosis (Thomas and Pommier, 2016; Rudin *et al*., 2021). It is marked by inactivation of the tumour suppressor genes *TP53* and *RB1* (George *et al*., 2015). Data from murine models and human tumors has suggested the existence of different SCLC subtypes with varying therapeutic vulnerabilities (Rudin *et al*., 2019; Simpson *et al*., 2020; Tlemsani *et al*., 2020; Gay *et al*., 2021). These efforts are reminiscent of earlier reports classifying SCLC as “classic” (neuroendocrine: NE) or “variant” (non-neuroendocrine: non-NE). Such intra-tumor phenotypic heterogeneity can aggravate disease progression by mechanisms such as cooperation among different phenotypes and evasion of many therapeutic assaults, as noted in SCLC and other cancers (Calbo *et al*., 2011; Chapman *et al*., 2014; Neelakantan *et al*., 2017; Jolly *et al*., 2018; Su *et al*., 2019). Given the distinct therapeutic vulnerabilities of the SCLC subtypes/phenotypes (Chen *et al*., 2021; Gay *et al*., 2021; Qu *et al*., 2021; Roper *et al*., 2021; Thomas *et al*., 2021; Yan *et al*., 2021), defining the molecular subtypes accurately can help rationalize novel therapeutic strategies for diverse patient cohorts.

Various efforts to characterize SCLC subtypes have identified different gene lists as classifiers. A 50-gene expression-based NE score was proposed to categorize SCLC tumors into NE and non-NE, which included 25 genes, each associated positively and negatively with NE differentiation status in cell lines (Zhang *et al*., 2018). Another list of 33 transcription factors (TFs) identified using NE and non-NE gene signatures suggested the possibility of a “hybrid” phenotype, implying that NE differentiation is not a strict “all-or-none” phenomenon (Udyavar *et al*., 2017). Starting from a cohort of SCLC samples, sets of co-expressed modules of genes were shortlisted and their upstream regulators were identified to arrive at these 33 TFs. These studies, and others, led to a classification of SCLC into four consensus phenotypes defined by enrichment of TFs: ASCL1 (SCLC-A), NEUROD1 (SCLC-N), YAP1 (SCLC-Y) and POU2F3 (SCLC-P) (Rudin *et al*., 2019). Another subtype – SCLC-A2 – driven by ASCL1 but different from SCLC-A has been proposed (Wooten *et al*., 2019), also supported by archetypal analysis of SCLC phenotypic landscape which proposed a 105-gene signature to classify phenotypes into five categories (Groves *et al*., 2021). Further, another SCLC subtype – marked not by enrichment of canonical TFs, but by enrichment of inflammation and immune-related genes (SCLC-I) – was proposed (Gay *et al*., 2021), further complicating ongoing attempts to arrive at a consensus of SCLC phenotypes and their distinct molecular and functional footprints.

Deciphering the regulatory networks driving phenotypic plasticity and heterogeneity and simulating their emergent dynamics has been helpful in mapping distinct phenotypes and corresponding molecular signatures in breast cancer and melanoma (Deshmukh *et al*., 2021; Pillai and Jolly, 2021). A similar approach applied to SCLC networks has identified stabilizers and destabilizers of various phenotypes: SCLC-A, SCLC-A2, and SCLC-N (Udyavar *et al*., 2017; Wooten *et al*., 2019), and revealed the underlying design principles of these networks (Chauhan *et al*., 2021). However, these SCLC networks did not include YAP1 and POU2F3, thus their ability to observe SCLC-Y and SCLC-P phenotypes was restricted. Moreover, how SCLC-I (characterized by low expression levels of ASCL1, NEUROD1 and POU2F3 (Gay *et al*., 2021)) emerges as a cell phenotype from the dynamics of these underlying networks, and the similarities and differences between SCLC-I and SCLC-Y phenotypes remains elusive.

Here, to identify similarities and differences in SCLC phenotypes proposed by different gene lists, we perform a comparative analysis across two SCLC cell line cohorts. Pairwise correlation among different gene lists reveal consistent behavior showing “teams” of different genes across gene lists both individually and in combination, suggesting some common molecular footprints of SCLC phenotypes identified across different studies. Moreover, we observed strongly positive correlation between SCLC-Y and SCLC-I phenotypes, indicating possible overlap between their programs. Finally, we report heterogeneity of SCLC phenotypes along the epithelial-mesenchymal transition (EMT) and NE axes. This work provides insights into sets of genes underlying such phenotypic heterogeneity and can pave a way toward reconciling seemingly divergent reports on mapping SCLC phenotypes.

## Results

### Diverse gene sets associated with SCLC phenotypic heterogeneity display consistent teams-like pattern

We calculated pairwise correlation coefficients for the set of 33 TFs (Udyavar *et al*., 2017) underlying phenotypic heterogeneity in SCLC. In both CCLE (n=49) and GSE73160 (n=67) (**Table S1**) cell line datasets, we observed that the pairwise correlation matrix revealed two sets of genes such that genes within a set were largely positively correlated with one another, but genes across these sets correlated negatively with one another. This striking observation indicated that 33 TFs can possibly form two “teams” of players such that players within a team activated each other but those across teams inhibited each other. Indeed, such “teams” like behavior was observed in a regulatory network identified for these 33 TFs (Chauhan et al., 2021; Udyavar et al., 2017). Intriguingly, one team comprised of NE genes such as *ASCL1* (Borromeo *et al*., 2016), *INSM1* (Fujino *et al*., 2015), *SOX2* (Voigt *et al*., 2021), *SOX11* (Walter *et al*., 2015), while the other one comprised genes corresponding to non-NE phenotype such as *MYC* (Mollaoglu *et al*., 2017; Ireland *et al*., 2020), *TEAD4* (Yan *et al*., 2021), *SMAD3* and *ZEB1* (Udyavar *et al*., 2017) This composition of teams raises the possibility that the two sets of genes forming a “toggle switch” (mutually repressive network) (Zhou and Huang, 2011) among each other corresponding to NE/non-NE differentiation in SCLC.

Next, we investigated whether this teams-like pattern was seen for other gene lists reported in the literature to categorize SCLC subtypes/phenotypic heterogeneity. For the 50-gene list used to calculate a NE score, we observed similar “teams” like pattern in pairwise correlation matrix for both CCLE and GSE73160 (red triangles and dark blue rectangles in **Fig 1A**). One of the two teams was composed of players associated with a NE phenotype – *ASCL1, INSM1, TAGNL3*, while the other team constituted players driving a non-NE phenotype – *YAP1* (Ito *et al*., 2016; Zhang *et al*., 2018). Similarly, for the 105 gene set, we discerned five such “teams” in both CCLE and GSE73160, thus pointing towards a generic theme for the existence of such mutually inhibiting teams corresponding to the specific SCLC phenotypes (**Fig 1B**). Combining the two gene lists (50 and 105 genes) maintained this “teams” like pattern in both the datasets. Addition of the 33 genes to this combined list continued to show similar trends, despite there being little overlap among the 33, 50 and 105 gene lists (**Fig S1; Table S2**). The consistency in patterns seen across individual and combinations of these independently identified gene lists point towards common molecular programs that operate towards enabling phenotypic heterogeneity in SCLC.

**Figure 1:**
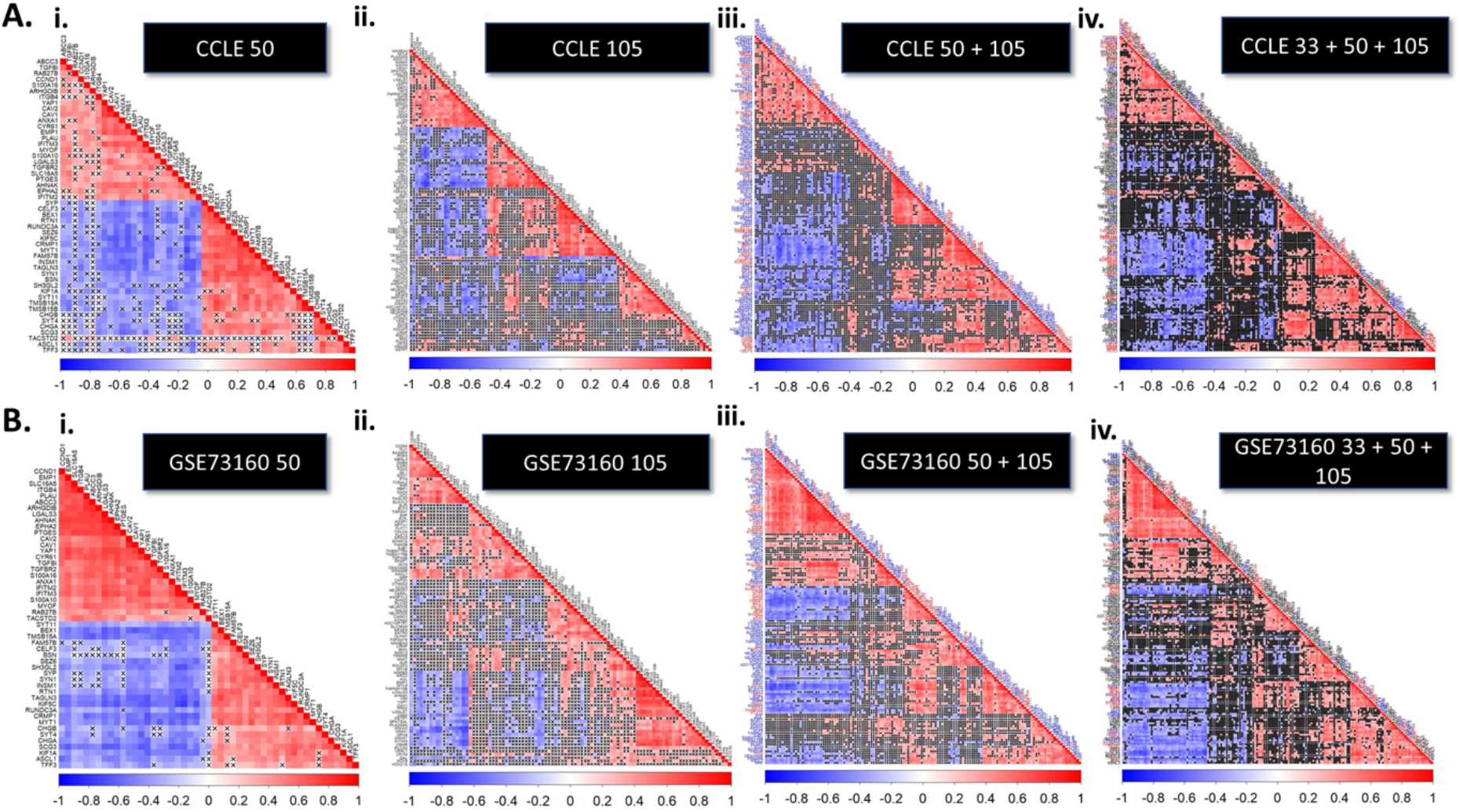
Existence of distinct groups of genes associated with phenotypic heterogeneity in SCLC. **A)** Matrices showing pairwise correlation coefficient (Spearman’s) of different markers of phenotypic heterogeneity for CCLE (n = 49) cell line dataset. i-iv represents different combinations of gene sets for SCLC signatures; i) 50 gene set; ii) 105 gene set; iii) 50 + 105 gene sets; iv) 33 + 50 + 105 gene sets. Crosses indicate p > 0.05. Colorbar denotes the values of correlation coefficient: red corresponds to positive correlation, blue for negative correlation. **B)** Same as A) but for GSE73160 (n = 67) cell line dataset.

### Presence of well-defined subtypes in SCLC

The 105 gene list comprised of markers for the 5 subtypes (SCLC-A, SCLC-A2, SCLC-N, SCLC-P and SCLC-Y) identified using archetypal analysis (Groves *et al*., 2021). Thus, we plotted the hierarchical clustering dendrogram to visualize five gene clusters prevalent in CCLE and GSE73160 datasets (**Fig 2A, i; 2B, i**). The gene-set clusters identified using these dendrograms had a high degree of similarity with the composition of groups reported earlier (Groves *et al*., 2021) (**Fig 2A,ii;Table S3**). The extent of overlap between the gene clusters identified in our study and the ones previously reported is represented as an overlap matrix. Each cell ranges from 0 to 1 and represents the fraction of genes in the previously reported clusters that are also present in the clusters identified in CCLE and GSE73160. These gene sets have been hereafter referred to as SCLC-A, SCLC-Y, SCLC-N, SCLC-P and based on presence of ASCL1, YAP1, NEUROD1, POU2F3, respectively. The 5^th^ gene cluster was referred to as SCLC-A2, based on the occurrence of genes reported earlier to be associated with SCLC-A2 subtype.

**Figure 2:**
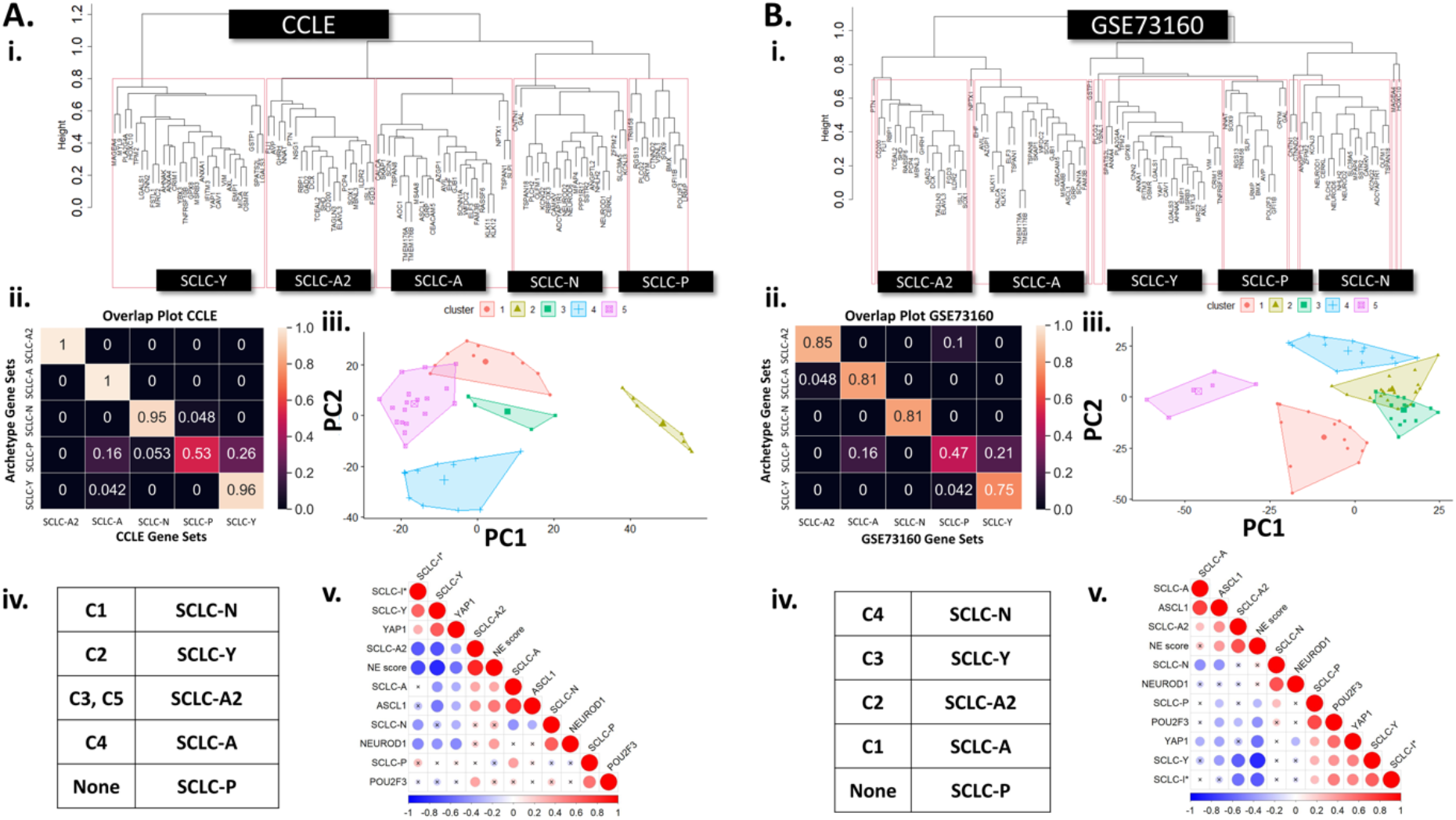
SCLC cell lines exhibit enrichment of distinct subtypes. **A)** i) Hierarchical clustering dendrograms showing presence of five different gene sets corresponding to the five SCLC subtypes in CCLE. Ii) Overlap of gene sets identified by dendrogram in panel i) and that by the archetypal analysis earlier. Colorbar represents the extent of overlap; numbers represent the fraction of genes corresponding to an archetype gene set which is also present in the said gene set identified via dendrogram iii) Principal Component Analysis (PCA) showing different clusters. iv) Table showing most enriched SCLC phenotype corresponding to each cluster identified by PCA. v) Pairwise correlation matrices (Spearman) for 11 metrics: ssGSEA scores of five SCLC subtypes identified by archetype analysis, that of SCLC I*, NE score and expression levels of four transcription factors (ASCL1, NEUROD1, POU2F3, YAP1). Crosses indicate p > 0.05. Colorbar denotes the values of correlation coefficient. **B)** Same as A) but for GSE73160.

Further, we asked whether these five gene sets identified from the dendrogram can help classify five proposed subtypes in SCLC. We used K-means clustering using K=5 for classifying five CCLE groups of samples. These five clusters segregate well when projected on their first two principal components (**Fig 2A, iii**). We next quantified the enrichment of the five gene sets identified via dendrogram for the different clusters of CCLE samples seen in CCLE, using single-sample gene set enrichment analysis (ssGSEA) (Barbie et al., 2009), to identify any possible one-to-one mapping between gene-sets and sample groups identified in CCLE. The cluster 1 seen in PCA plot (**Fig 2A, iii**) was enriched for SCLC-N geneset; cluster 2 was enriched for SCLC-Y geneset; cluster 4 was enriched for SCLC-A, as compared to other gene sets (**Fig S2A**). Clusters 3 and 5 showed comparable enrichment of SCLC-A2, and all clusters had comparable enrichment of SCLC-P geneset; thus, the clusters 3 and 5 can be mapped to SCLC-A2 geneset, while SCLC-P corresponded to a mixture of samples from diverse groups seen in the PCA plot (**Fig 2A, iv**). Thus, the ssGSEA analysis resulted in the assignment of these five clusters to four unique gene sets.

Further, we plotted a pairwise correlation matrix for 11 metrics (**Table S4**) comprising of a) ssGSEA scores of five gene sets as identified from archetypal analysis (Groves *et al*., 2021), b) expression levels of four TFs associated with phenotypic heterogeneity in SCLC – ASCL1, YAP1, NEUROD1 and, POU2F3, c) NE score (Zhang *et al*., 2018) (the higher the NE score, the more NE phenotype; the lower the NE score, the more non-NE phenotype), and d) ssGSEA score of the SCLC-I* gene set representing a phenotype similar to the one (SCLC-inflamed or SCLC-I) reported recently showing enrichment of inflamed gene signature (Gay *et al*., 2021). Interestingly, we observed that eight metrics (NE scores, ssGSEA scores of SCLC-A, SCLC-N, SCLC-A2 and SCLC-P, expression levels of ASCL1, NEUROD1 and POU2F3) positively correlated with one another, while the other three metrics (ssGSEA scores of SCLC-Y, SCLC-I* and expression levels of YAP1) positively correlated with one another (**Fig 2A, v**). These eight metrics were largely negatively correlated with the other three, reminiscent of “teams”-like behavior as noted earlier for gene expression based correlation matrices. Similar classification was evident in other distance-based dendrograms for these 11 metrics as well as for the ssGSEA scores of the 5 archetype based gene-sets (**Fig S3**), indicating that SCLC-Y was sufficiently different from other four SCLC subtypes (SCLC-A, SCLC-P, SCLC-A2 and SCLC-N). Also, SCLC-Y was the only subtype whose enrichment correlated negatively with NE score. Largely consistent patterns were observed in GSE73160 as well (**Fig 2B,ii-v; S3**), pinpointing that five well-defined subtypes of SCLC exist spread along the NE/non-NE spectrum.

### Similarity between transcriptional signatures of SCLC-I* and SCLC-Y subtypes

Intrigued by the observations, in both CCLE and GSE73160, that a) the SCLC-I* (**Table S5**) enrichment was positively correlated with that of SCLC-Y and that b) SCLC-I* and SCLC-Y enrichment correlated negatively with NE scores, we examined whether SCLC-I* ssGSEA scores were negatively correlated with NE scores in additional independent datasets. First, we observed that the set of cell lines in CCLE and GSE73160 that were associated with the lowest NE scores (cluster 2 in CCLE, and cluster 3 in GSE73160, both mapping to SCLC-Y respectively) had the highest ssGSEA scores for SCLC-I* subtype (**Fig S4**). Next, in CCLE, GSE73160 as well as in six additional SCLC cell line datasets, we observed consistent patterns showing negative correlation between the NE and the SCLC-I* enrichment scores (**Fig 3A, B; Fig S5**), thus confirming a non-NE status of SCLC-inflamed subtype. Thus, put together, the correlation matrices and enrichment scores for different sets of cell lines in CCLE and GSE73160 indicates a high degree of overlap in molecular and/or functional footprints of the SCLC-Y and SCLC-I* subtypes.

**Figure 3:**
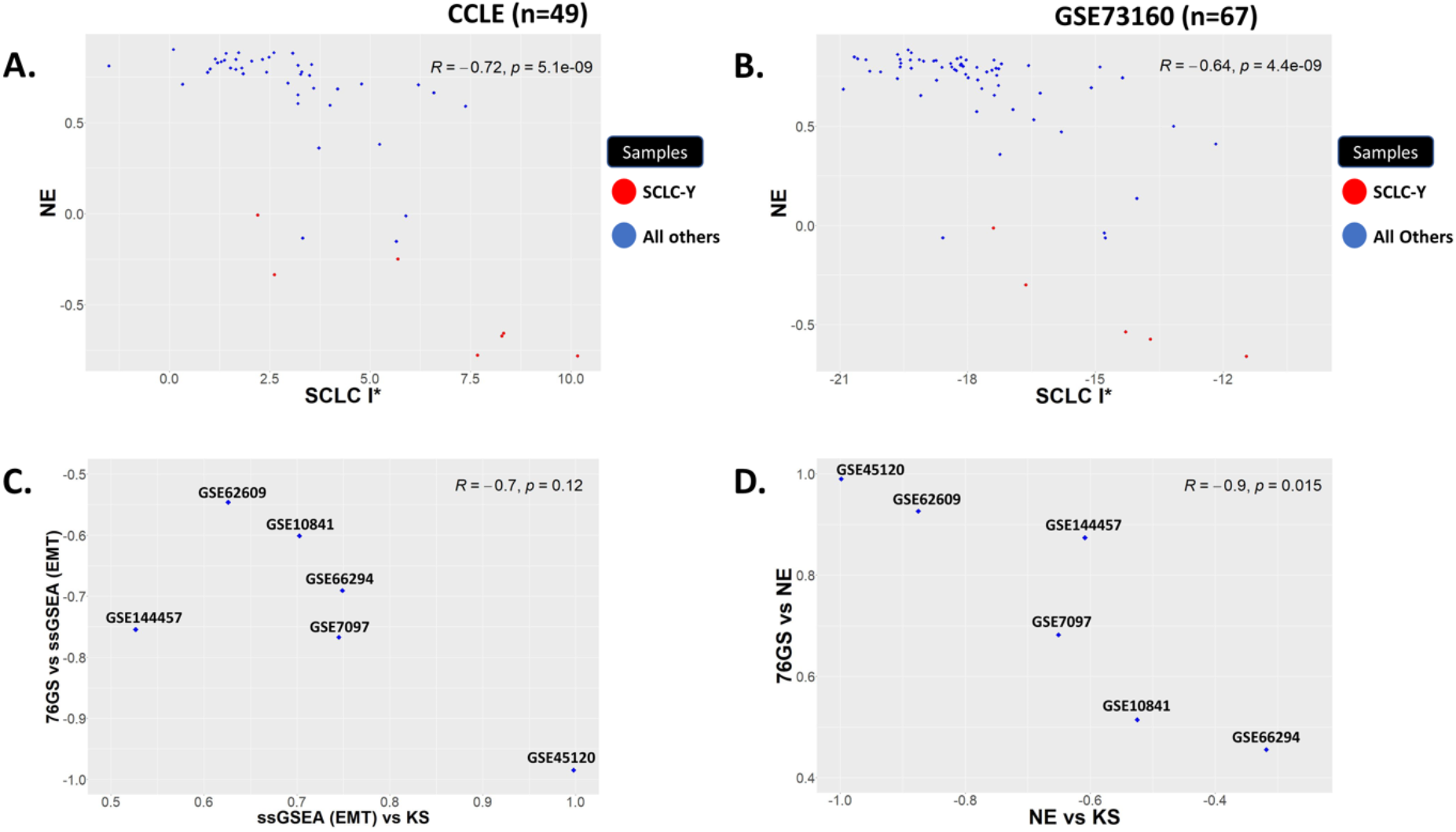
EMT, NE scores, and YAP1 levels in independent SCLC datasets. **A, B)** Scatter plots showing the correlation between NE scores (Y-axis) and SCLC I* ssGSEA scores (X-axis) for CCLE (n=49) and GSE73160 (n=67). **C)** Scatter plot showing the Pearson correlation coefficients of two different EMT scores (76GS and KS) with ssGSEA scores for Hallmark MSigDB EMT gene set. Each dot represents a different SCLC cell line dataset (GSE ID indicated) for which these correlation coefficients have been calculated. **D)** Same as C) but for correlation of NE scores with 76GS and KS EMT scores.

NE/non-NE subtypes in SCLC have also been proposed to have varying EMT status (Udyavar *et al*., 2017); thus, we quantified the extent of EMT in SCLC cell lines using three different metrics: 76GS, KS and ssGSEA scores for MSigDB EMT signature. The higher the 76GS score, the more epithelial the sample is. The higher the KS or the ssGSEA (EMT) score, the more mesenchymal the sample is. Thus, we expect KS and ssGSEA (EMT) scores to be positively correlated with one another, while 76GS to be negatively correlated with these two metrics, as seen for other cancers (Chakraborty *et al*., 2020; Mandal *et al*., 2021). Indeed, across the six additional SCLC cell line datasets, we notice expected trends, thus revealing consistency in these metrics to quantify EMT (**Fig 3C**). Interestingly, for these datasets, the NE scores were positively correlated with 76GS scores but negatively with KS scores (**Fig 3D**), thus suggesting that SCLC cell lines where EMT markers are expressed tend to be less neuroendocrine (NE). Given the association of EMT with immune evasion (Tripathi *et al*., 2016; Sahoo *et al*., 2021), enrichment of inflamed gene signature, as well as mesenchymal signature in non-NE cell lines (**Table S6**), suggests a convergence of non-neuroendocrine behavior, YAP1 levels, SCLC inflamed subtype, and mesenchymal features.

## Discussion

Phenotypic (non-genetic) heterogeneity is emerging as a major driver of metastasis and therapy recalcitrance across multiple cancers. While genetic heterogeneity has been well-expounded, the causes and consequences of phenotypic heterogeneity are relatively less explored (Bell and Gilan, 2020; Marine *et al*., 2020). Phenotypic heterogeneity in SCLC has been investigated since the 1980s when ‘adherent’ and ‘variant’ cells were reported, but the use of different gene lists and functional assays and relatively smaller sample size has obfuscated the identification of molecular and functional attributes of various subtypes/phenotypes as well as their actionable therapeutic vulnerabilities. A consensus classification for SCLC has been proposed, where a tumor sample could be classified into SCLC-N, SCLC-A, SCLC-P and SCLC-Y subtypes based on the relative expression levels of TFs NEUROD1, ASCL1, POU2F3 and YAP1 (Rudin *et al*., 2019). However, a recent study identified an additional subtype not marked by enrichment of any of these 4 TFs, but by that of inflammation and immune-evasive signatures, thus termed SCLC-I (Gay *et al*., 2021). Similarly, SCLC-A2/NEv2 phenotype has been reported as well with a distinct drug-resistance signature (Wooten *et al*., 2019). Together, these classifications raise the questions: a) how many SCLC subtypes exist preclinically/ clinically? b) what is the molecular footprint of each subtype?, c) which subtype is the most aggressive one? and d) what are the subtype-specific vulnerabilities? Here, we demonstrated that despite little or no overlap in terms of the individual genes, there exist “teams” of co-expressed modules of genes that capture the phenotypic heterogeneity of distinct SCLC subtypes. Such “teams” may comprise of drivers of NE/non-NE phenotype as well as additional players enabling heterogeneity within this broader classification. Such ‘teams’ have been observed between EMT-inducers and EMT-inhibitors (Jia *et al*., 2020) and between drivers of proliferative and invasive phenotypes in melanoma (Pillai and Jolly, 2021), indicating a potential common design principle for the networks involved in cancer cell plasticity. We also demonstrated a high degree of transcriptional similarity between the subtypes negatively correlated with high NE scoring: SCLC-Y and SCLC-I*, reminiscent of recent observations that YAP1 expression in SCLC describes a T-cell inflamed subtype (Owonikoko *et al*., 2021). SCLC-A and SCLC-N enrichment was also positively correlated in CCLE and GSE73160, but they are regarded as distinct subtypes at a community consensus level. Whether SCLC-I and SCLC-Y are two distinct programs or have largely overlapping arms needs to be investigated further.

Our analysis so far includes only two cell line datasets. To what extent are these trends maintained in patient tumors at a bulk and individual cell level remains to be seen. Further, our work so far does not offer insights into the dynamics of cell-state transition from a mechanistic standpoint. Nonetheless, these preliminary trends suggest carefully revisiting the classification of SCLC-I and SCLC-Y as different subtypes, and mapping out their similarities and differences at molecular and functional levels. Moreover, connecting the metastatic potential and therapy evasion to subtypes and their EMT status (Yan *et al*., 2021) would lead to a better understanding of underlying networks driving reversible phenotypic plasticity and heterogeneity in SCLC.

## Materials and Methods

### RNA-seq & microarray data

We used 50 Small Cell Lung Cancer (SCLC) cell lines from the Cancer Cell Line Encyclopedia (CCLE) (Barretina *et al*., 2012) dataset, and GSE73160 (Polley *et al*., 2016) (n=67 SCLC cell lines from the National Cancer Institute). For additional analysis, we used GSE7097 (Olejniczak *et al*., 2007), GSE10841 (Tse *et al*., 2008), GSE66294 (Mohammad *et al*., 2015), GSE45120 (Canadas *et al*., 2014), GSE62609 (Christensen *et al*., 2014), GSE144457 (Llabata *et al*., 2021) and GSE161740 (Wu *et al*., 2021). All analyses were carried out using R version 4.0.4 and Python version 3.8.3 unless mentioned otherwise. The SCLC CCLE RNA-seq dataset was Transcripts Per Million (TPM) normalized. From the probe-wise expression matrix, all microarray datasets were pre-processed to provide gene-wise expression for each sample. In case of multiple probes mapping to a single gene, median gene expression of all mapped probes was considered for that gene. *bioMart* package from Bioconductor version 3.11 was used for mapping of probes to gene ids across different platforms (Entrez id, Ensembl id or Affymetrix probe id).

### Correlation plots

We determined the extent of correlation between a given pair of genes using Pearson’s correlation coefficient and Spearman’s rank correlation coefficient. Corresponding correlation coefficient values vary between −1 to +1. +1 indicating a strong positive correlation and −1 indicating a strongly negative one. R package *corrplot* was used to plot the correlation matrices.

### Identification of gene sets for individual subtypes

Hierarchical clustering was performed using the *hclust* function from the *fastcluster* package with 1-corr as the dissimilarity metric, where corr is the Spearman correlation matrix for the subset of genes or the gene sets in a particular dataset. The *cutree* function from the *stats* package was used to cut the generated dendrogram at a specified height or into the desired number of groups or gene clusters.

### Principal Component Analysis (PCA)

To visualize multidimensional RNA-seq and microarray expression datasets, we used PCA. R package *factoextra* was used to calculate and visualize the data points on the principal components.

### Identification of SCLC subtypes

SCLC-A, SCLC-A2, SCLC-N, SCLC-P and SCLC-Y subtypes were mapped to the corresponding clusters generated by k-means using the Single-sample GSEA (ssGSEA) (Barbie *et al*., 2009). ssGSEA is an extension of Gene Set Enrichment Analysis (GSEA) (Subramanian *et al*., 2005) and it calculates separate enrichment scores for each pair of a sample and a gene set. ssGSEA enrichment score quantifies the degree to which the genes in a particular gene set are coordinately up-regulated or down-regulated within each sample.

### Inflamed signature (SCLC-I*) calculation

The non-negative matrix factorization (NMF) defined gene list having 1300 genes was used as a baseline gene set (Gay et al., 2021). In this study, SCLC-inflamed or SCLC-I subtype was found to uniquely express numerous immune checkpoint genes and human leukocyte antigens (HLAs). Thus, we downloaded various targetable immune checkpoints and their ligands from the abovementioned study as well as similar gene sets from the Molecular Signatures Database (MSigDB) (**Table S5**) (Liberzon *et al*., 2011) (<u>GNF2_HLA_C</u>, <u>IMMUNE_RESPONSE</u>, <u>IMMUNE_SYSTEM_PROCESS</u>, <u>GOMF_CHEMOKINE__ACTIVITY</u>, <u>REACTOME_INNATE_IMMUNE_SYSTEM</u>, <u>REACTOME_ADAPTIVE_IMMUNE_SYSTEM</u>). The intersection of MSigDB gene sets and 1300 gene set was used for ssGSEA analysis for individual datasets. We generated ssGSEA scores for SCLC-I* subtype to be used for downstream analysis.

### NE score calculation

NE scores were calculated as previously reported using 50 genes. This list was identified by using the most highly differentially expressed genes (25 genes overexpressed in NE cell lines, 25 genes overexpressed in non-NE cell lines) (Zhang et al., 2018). From this signature, a quantitative NE score is generated using the formula: NE score = (correl NE – correl non-NE)/2. Here correl NE (or non-NE) is the Pearson correlation between expression of 50 genes in the test sample and expression of these genes in neuroendocrine (NE) or non-NE (non-neuroendocrine) cell line group. The NE score ranges from −1 to +1, where a positive score predicts NE and an epithelial phenotype while a negative score predicts non-NE cell types.

### Calculation of EMT Scores

EMT scores (76GS and KS) were calculated as reported earlier (Chakraborty *et al*., 2020). 76GS scores were calculated using a 76-gene expression signature and a metric based on that gene signature (Byers *et al*., 2013; Guo *et al*., 2019). For each sample, we calculated the score as a weighted sum of 76 gene expression levels, with the weight being the value of correlation coefficient of expression levels of *CDH1* and the corresponding gene. Negative (resp. positive) scores correspond to Mesenchymal (M) (resp. epithelial (E)) phenotype. KS EMT scores were calculated according to the procedure reported earlier based on difference of cumulative distribution functions (CDFs) of E and M signatures in a dataset. M samples have a positive KS score, whereas E samples have a negative KS score (Tan *et al*., 2014).

## Author contributions

MKJ and AT designed and supervised research; AS, PD, MP, NA and NT performed research and analysed data. All authors contributed to writing and editing of manuscript.

## Data and code availability

All transcriptomic data used in the analysis is publicly available. All codes are available at https://github.com/abhay8154/SCLC-Project-Code

## Supplementary Figures

**Figure S1:**
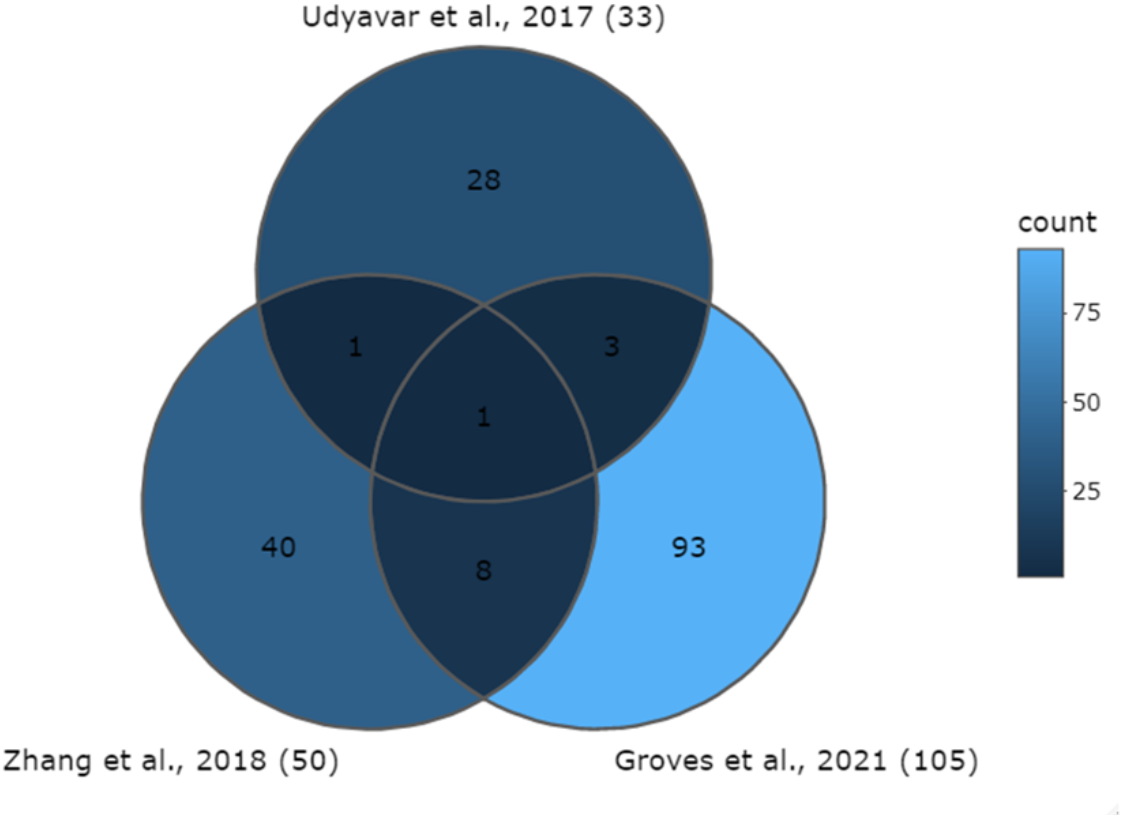
Venn diagram showing overlap of Groves et al (105 genes by Archetypal Analysis), Zhang et al (50 genes) and Udyavar et al (33 genes) gene sets

**Figure S2:**
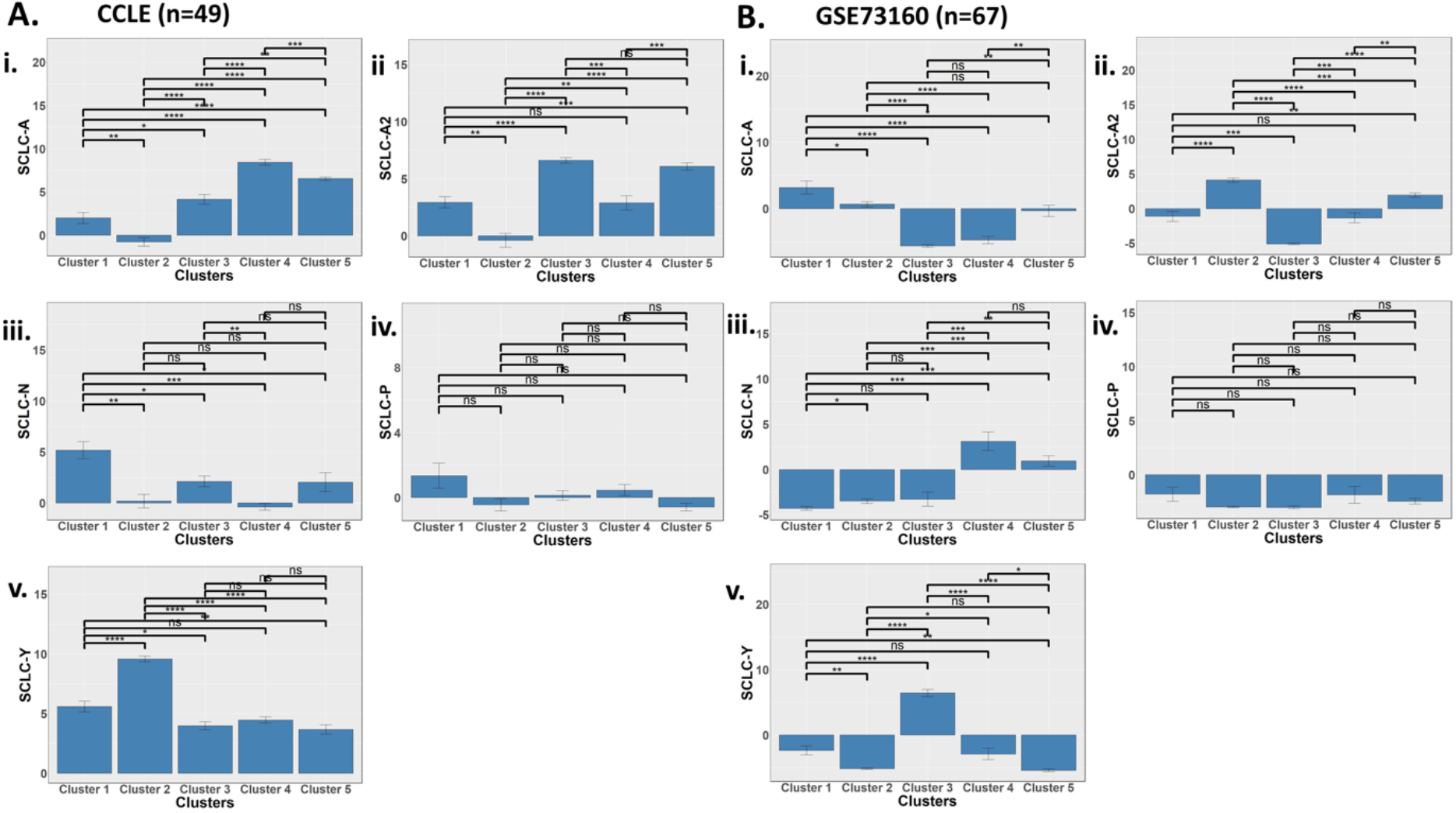
Bar plots showing cluster wise ssGSEA enrichment scores for five Archetypal analysis gene sets i. SCLC-A, ii. SCLC-A2, iii. SCLC-N, iv. SCLC-P, v. SCLC-Y for A) CCLE, B) GSE73160 cell lines. 2. (Significance is represented based on FDR adjusted p-value for two-sided Welch’s t-test, * for p < 0.05, ** for p < 0.01, *** for p < 0.001, **** for p < 0.0001 and ns for p ≥ 0.05 i.e., nonsignificant).

**Figure S3:**
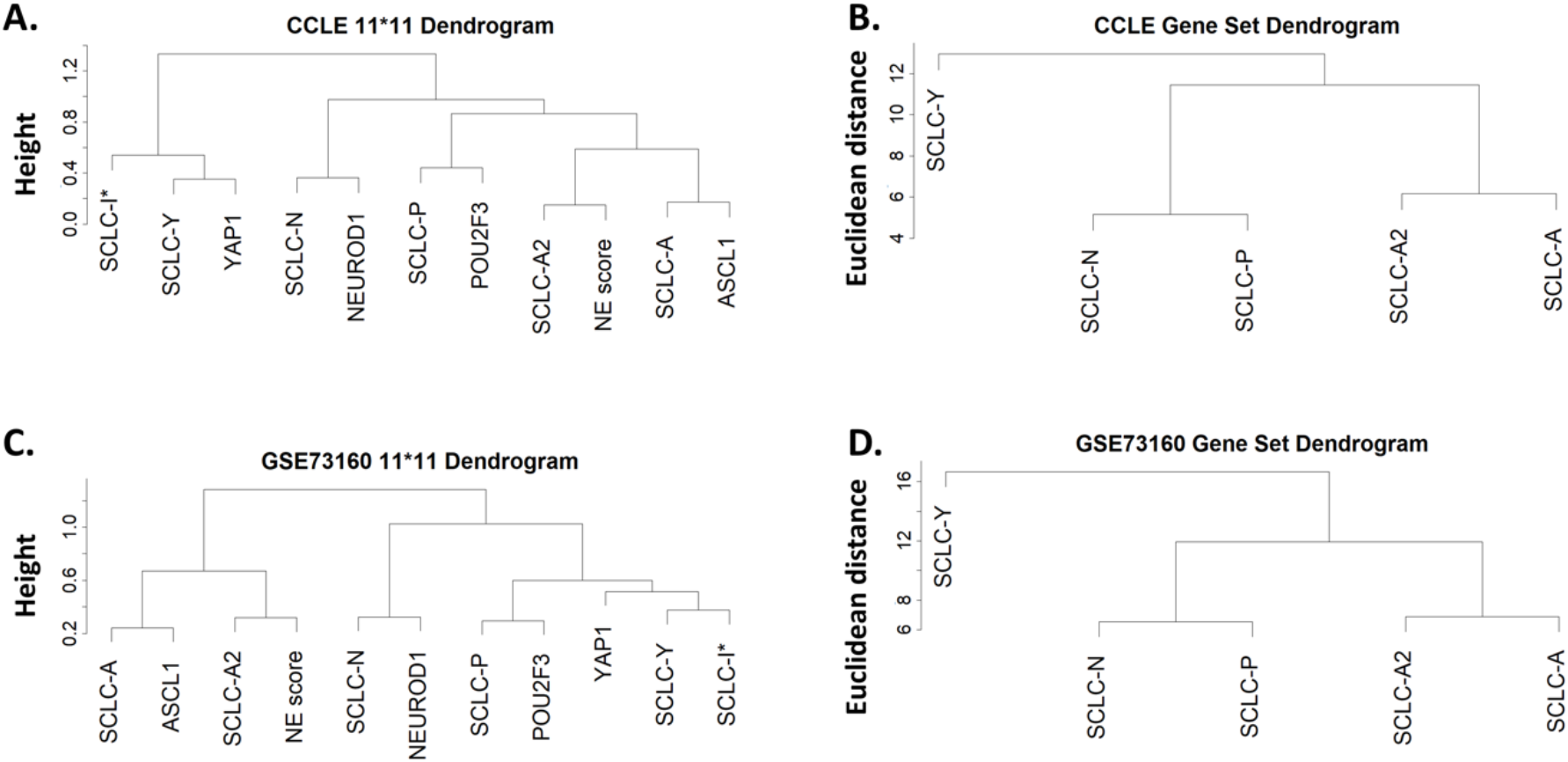
A, C) Hierarchical Clustering dendrograms showing the 11*11 correlation matrix plotted in Fig. 2B, 2G. SCLC-A, SCLC-A2, SCLC-N, SCLC-P, SCLC-Y and SCLC-I* denote ssGSEA scores for all the samples in individual datasets. ASCL1, NEUROD1, POU2F3 and YAP1 denote the mean expression levels of respective transcription factors across all the samples in CCLE (S3A) and GSE63170 (S3C). B, D) Hierarchical clustering dendrograms showing the relationship among different Archetypal Analysis gene sets for CCLE (S3B) and GSE73160 (S3D).

**Figure S4:**
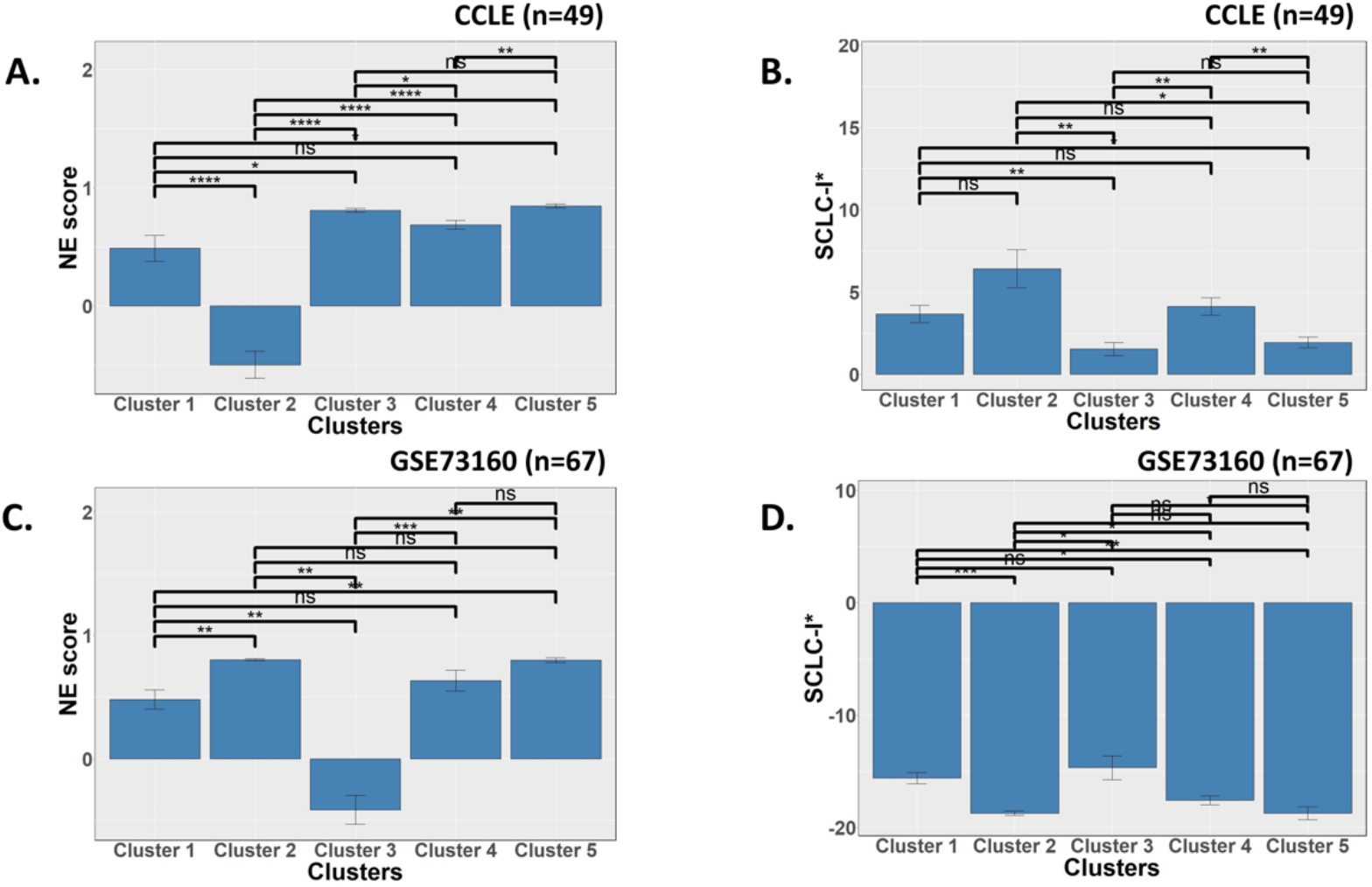
A, C) Bar plots showing cluster wise NE score for CCLE (S4A) and GSE73160 (S4C). B, D) Bar plots showing cluster wise ssGSEA scores for SCLC-I* (Inflamed subtype) for CCLE (S4B) and GSE73160 (S4D). Statistical significance is represented based on FDR adjusted p-value for two-sided Welch’s t-test, * for p < 0.05, ** for p < 0.01, *** for p < 0.001, **** for p < 0.0001 and ns for p ≥ 0.05 i.e., non-significant.

**Fig S5:**
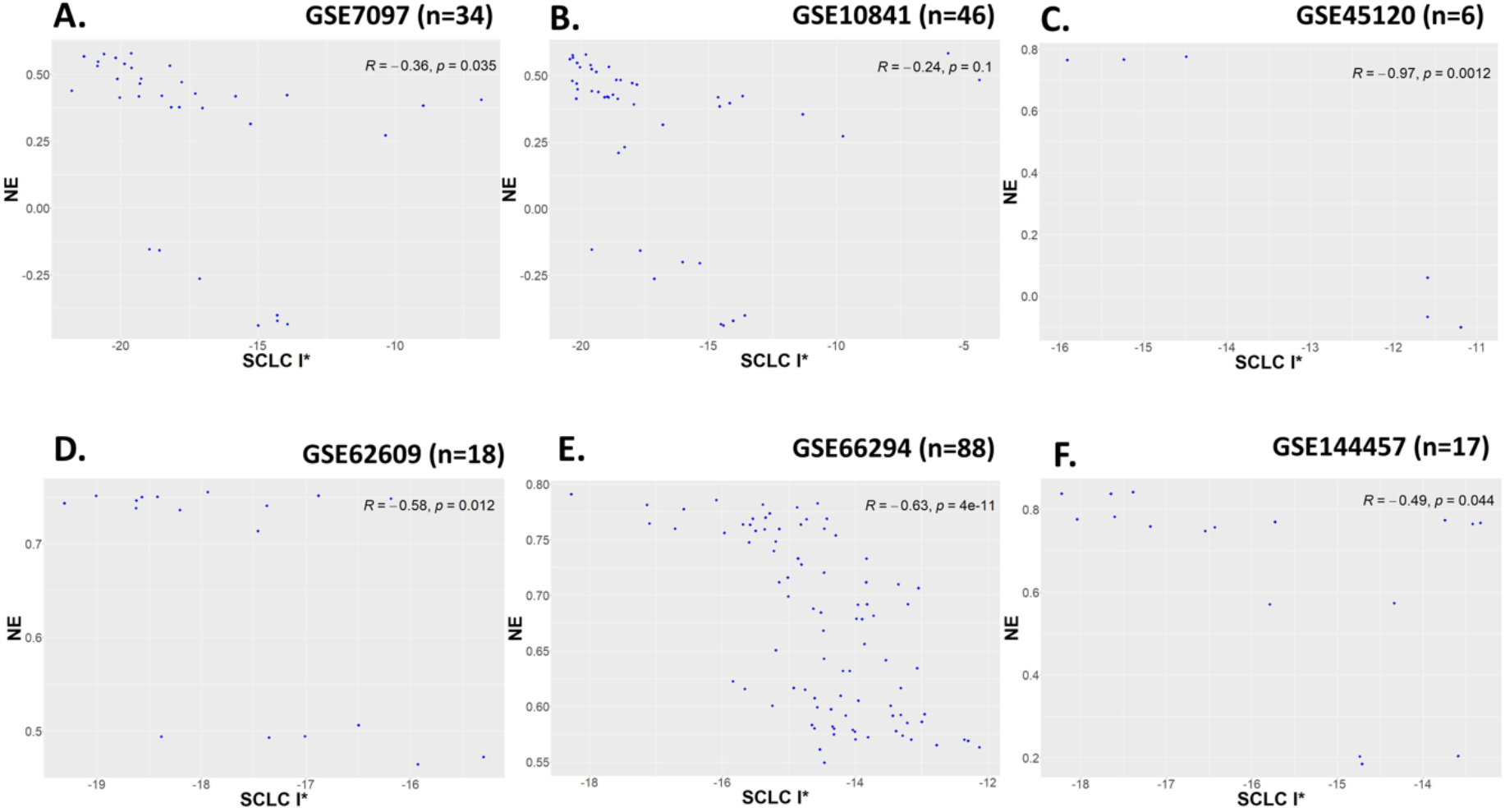
Scatter plots showing correlation between NE scores (y-axis) and SCLC I* ssGSEA scores (x-axis) for six additional datasets: A-F) GSE7097 (n=34), GSE10841 (n=46), GSE45120 (n=6), GSE62609 (n=18), GSE66294 (n=88) and GSE144457 (n=17) SCLC microarray cell line datasets. R, p denotes the Pearson correlation coefficient and corresponding p-value respectively.

## Supplementary Table Legends

**Table S1:** List of cell lines in CCLE and GSE73160

**Table S2**: Composition of teams seen using 105 genes, 50 genes, 105 genes+ 50 genes, and 105 genes+50 genes + 33genes for CCLE and GSE73160

**Table S3:** Clusters seen from dendrogram of 105 genes for CCLE and GSE73160

**Table S4:** Correlation coefficient values seen for 11*11 pairwise correlation for CCLE and GSE73160

**Table S5:** Gene list used for SCLC-I* for different datasets

**Table S6**: NE, SCLC-I* ssGSEA, KS and 76GS scores for different datasets

## References

Barbie, DA et al. (2009). Systematic RNA interference reveals that oncogenic KRAS-driven cancers require TBK1. Nature 462, 108–112.

Barretina, J et al. (2012). The Cancer Cell Line Encyclopedia enables predictive modelling of anticancer drug sensitivity. Nature 483, 603–607.

Bell, CC, and Gilan, O (2020). Principles and mechanisms of non-genetic resistance in cancer. Br J Cancer 122, 465–472.

Borromeo, MD et al. (2016). ASCL1 and NEUROD1 Reveal Heterogeneity in Pulmonary Neuroendocrine Tumors and Regulate Distinct Genetic Programs. Cell Rep 16, 1259–1272.

Byers, LA et al. (2013). An epithelial-mesenchymal transition gene signature predicts resistance to EGFR and PI3K inhibitors and identifies Axl as a therapeutic target for overcoming EGFR inhibitor resistance. Clin Cancer Res 19, 279–290.

Calbo, J, van Montfort, E, Proost, N, van Drunen, E, Beverloo, HB, Meuwissen, R, and Berns, A (2011). A functional role for tumor cell heterogeneity in a mouse model of small cell lung cancer. Cancer Cell 19, 244–256.

Canadas, I et al. (2014). Targeting epithelial-to-mesenchymal transition with met inhibitors reverts chemoresistance in small cell lung cancer. Clin Cancer Res 20, 938–950.

Chakraborty, P, George, JT, Tripathi, S, Levine, H, and Jolly, MK (2020). Comparative Study of Transcriptomics-Based Scoring Metrics for the Epithelial-Hybrid-Mesenchymal Spectrum. Front Bioeng Biotechnol 8, 220.

Chapman, A, del Ama, LF, Ferguson, J, Kamarashev, J, Wellbrock, C, and Hurlstone, A (2014). Heterogeneous tumor subpopulations cooperate to drive invasion. Cell Rep 8, P688–695.

Chauhan, L, Ram, U, Hari, K, and Jolly, MK (2021). Topological signatures in regulatory network enable phenotypic heterogeneity in small cell lung cancer. Elife 10, e64522.

Chen, H, Gesumaria, L, Park, Y-K, Oliver, TG, Singer, DS, Ge, K, and Shrump, DS (2021). BET Bromodomain Inhibitors Target the NEUROD1-subtype SCLC by Blocking NEUROD1 Transactivation. BioRxiv, 4665771.

Christensen, CL et al. (2014). Targeting Transcriptional Addictions in Small Cell Lung Cancer with a Covalent CDK7 Inhibitor. Cancer Cell 26, 909–922.

Deshmukh, AP et al. (2021). Identification of EMT signaling cross-talk and gene regulatory networks by single-cell RNA sequencing. Proc Natl Acad Sci U S A 118, e2102050118.

Fujino, K et al. (2015). Insulinoma-associated protein 1 is a crucial regulator of neuroendocrine differentiation in lung cancer. Am J Pathol 185, 3164–3177.

Gay, CM et al. (2021). Patterns of transcription factor programs and immune pathway activation define four major subtypes of SCLC with distinct therapeutic vulnerabilities. Cancer Cell 39, 346–360.E7.

Gazdar, AF, Bunn, PA, and Minna, JD (2017). Small-cell lung cancer: What we know, what we need to know and the path forward. Nat Rev Cancer 17, 725–737.

George, J et al. (2015). Comprehensive genomic profiles of small cell lung cancer. Nature 524, 47–53.

Groves, SM, Ireland, A, Liu, Q, Simmons, AJ, Lau, K, Iams, WT, Tyson, D, Lovly, CM, Oliver, TG, and Quaranta, V (2021). Cancer Hallmarks Define a Continuum of Plastic Cell States between Small Cell Lung Cancer Archetypes. BioRxiv, 427865.

Guo, CC et al. (2019). Dysregulation of EMT Drives the Progression to Clinically Aggressive Sarcomatoid Bladder Cancer. Cell Rep 27, 1781–1793.e4.

Ireland, AS et al. (2020). MYC Drives Temporal Evolution of Small Cell Lung Cancer Subtypes by Reprogramming Neuroendocrine Fate. Cancer Cell 38, 60–78.e12.

Ito, T et al. (2016). Loss of YAP1 defines neuroendocrine differentiation of lung tumors. Cancer Sci 107, 1537–1538.

Jia, W, Tripathi, S, Chakraborty, P, Chedere, A, Rangarajan, A, Levine, H, and Jolly, MK (2020). Epigenetic feedback and stochastic partitioning during cell division can drive resistance to EMT. Oncotarget 11, 2611–2624.

Jolly, MK, Kulkarni, P, Weninger, K, Orban, J, and Levine, H (2018). Phenotypic Plasticity, Bet-Hedging, and Androgen Independence in Prostate Cancer: Role of Non-Genetic Heterogeneity. Front Oncol 8, 50.

Liberzon, A, Subramanian, A, Pinchback, R, Thorvaldsdóttir, H, Tamayo, P, and Mesirov, JP (2011). Molecular signatures database (MSigDB) 3.0. Bioinformatics 27, 1739–1740.

Lin, CA, Yu, SL, Chen, HY, Chen, HW, Lin, SU, Chang, CC, Yu, CJ, Yang, PC, and Ho, CC (2019). EGFR-Mutant SCLC Exhibits Heterogeneous Phenotypes and Resistance to Common Antineoplastic Drugs. J Thorac Oncol 14, 513–526.

Llabata, P et al. (2021). MAX mutant small-cell lung cancers exhibit impaired activities of MGA-dependent noncanonical polycomb repressive complex. Proc Natl Acad Sci U S A 118, e2024824118.

Mandal, S et al. (2021). Transcriptomic-based quantification of the epithelial-hybrid-mesenchymal spectrum across biological contexts. BioRxiv, 458982.

Marine, J-C, Dawson, S-J, and Dawson, MA (2020). Non-genetic mechanisms of therapeutic resistance in cancer. Nat Rev Cancer 20, 743–756.

Mohammad, HP et al. (2015). A DNA Hypomethylation Signature Predicts Antitumor Activity of LSD1 Inhibitors in SCLC. Cancer Cell 28, 57–69.

Mollaoglu, G et al. (2017). MYC Drives Progression of Small Cell Lung Cancer to a Variant Neuroendocrine Subtype with Vulnerability to Aurora Kinase Inhibition. Cancer Cell 31, 270–285.

Neelakantan, D et al. (2017). EMT cells increase breast cancer metastasis via paracrine GLI activation in neighbouring tumour cells. Nat Commun 8, 15773.

Olejniczak, ET, Van Sant, C, Anderson, MG, Wang, G, Tahir, SK, Sauter, G, Lesniewski, R, and Semizarov, D (2007). Integrative genomic analysis of small-cell lung carcinoma reveals correlates of sensitivity to Bcl-2 antagonists and uncovers novel chromosomal gains. Mol Cancer Res 5, 331–339.

Owonikoko, TK et al. (2021). YAP1 Expression in SCLC Defines a Distinct Subtype With T-cell– Inflamed Phenotype. J Thorac Oncol 16, 464–476.

Pillai, M, and Jolly, MK (2021). Systems-level network modeling deciphers the master regulators of phenotypic plasticity and heterogeneity in melanoma. IScience 24, 103111.

Polley, E et al. (2016). Small Cell Lung Cancer Screen of Oncology Drugs, Investigational Agents, and Gene and microRNA Expression. J Natl Cancer Inst 108, djw122.

Qu, S, Fetsch, P, Thomas, A, Pommier, Y, Schrump, DS, Miettinen, MM, and Chen, H (2021). Molecular Subtypes of Primary SCLC Tumors and Their Associations With Neuroendocrine and Therapeutic Markers. J Thorac Oncol, in press.

Roper, N et al. (2021). Notch signaling and efficacy of PD-1/PD-L1 blockade in relapsed small cell lung cancer. Nat Commun 12, 3880.

Rudin, CM et al. (2019). Molecular subtypes of small cell lung cancer: a synthesis of human and mouse model data. Nat Rev Cancer 19, 289–297.

Rudin, CM, Brambilla, E, Faivre-Finn, C, and Sage, J (2021). Small-cell lung cancer. Nat Rev Dis Prim 7, 3.

Sahoo, S, Nayak, SP, Hari, K, Purkait, P, Mandal, S, Kishore, A, Levine, H, and Jolly, MK (2021). Immunosuppressive traits of the hybrid epithelial/mesenchymal phenotype. BioRxiv, 449285.

Simpson, KL et al. (2020). A biobank of small cell lung cancer CDX models elucidates inter- and intratumoral phenotypic heterogeneity. Nat Cancer 1, 437–451.

Su, Y, Bintz, M, Yang, Y, Robert, L, Ng, AHC, Liud, V, Ribas, A, Heath, JR, and Wei, W (2019). Phenotypic heterogeneity and evolution of melanoma cells associated with targeted therapy resistance. PLoS Comput Biol 15, e1007034.

Subramanian, A et al. (2005). Gene set enrichment analysis: A knowledge-based approach for interpreting genome-wide expression profiles. Proc Natl Acad Sci U S A 102, 15545–15550.

Tan, TZ, Miow, QH, Miki, Y, Noda, T, Mori, S, Huang, RY, and Thiery, JP (2014). Epithelial-mesenchymal transition spectrum quantification and its efficacy in deciphering survival and drug responses of cancer patients. EMBO Mol Med 6, 1279–1293.

Thomas, A et al. (2021). Therapeutic targeting of ATR yields durable regressions in small cell lung cancers with high replication stress. Cancer Cell 39, 566–579.e7.

Thomas, A, and Pommier, Y (2016). Small cell lung cancer: Time to revisit DNA-damaging chemotherapy. Sci Transl Med 8, 346fs12.

Tlemsani, C et al. (2020). SCLC-CellMiner : A Resource for Small Cell Lung Cancer Cell Line Genomics and Pharmacology Based on Genomic Signatures ll SCLC-CellMiner : A Resource for Small Cell Lung Cancer Cell Line Genomics and Pharmacology Based on Genomic Signatures. Cell Rep 33, 108296.

Tripathi, SC et al. (2016). Immunoproteasome deficiency is a feature of non-small cell lung cancer with a mesenchymal phenotype and is associated with a poor outcome. Proc Natl Acad Sci U S A 113, E1555–64.

Tse, C et al. (2008). ABT-263: A potent and orally bioavailable Bcl-2 family inhibitor. Cancer Res 68, 3421–3428.

Udyavar, AA, Wooten, DJ, Hoeksema, M, Bansal, M, Califano, M, Estrada, L, Schnell, S, Irish, JM, Massion, PP, and Quaranta, V (2017). Novel Hybrid Phenotype Revealed in Small Cell Lung Cancer by a Transcription Factor Network Model That Can Explain Tumor Heterogeneity. Cancer Res 77, 1063–1074.

Voigt, E et al. (2021). Sox2 Is an Oncogenic Driver of Small-Cell Lung Cancer and Promotes the Classic Neuroendocrine Subtype. Mol Cancer Res, in press.

Walter, RFH et al. (2015). SOX4, SOX11 and PAX6 mRNA expression was identified as a (prognostic) marker for the aggressiveness of neuroendocrine tumors of the lung by using nextgeneration expression analysis (NanoString). Futur Med 11, 1027–1036.

Wooten, DJ, Groves, SM, Tyson, DR, Liu, Q, Lim, JS, Albert, R, Lopez, CF, Sage, J, and Quaranta, V (2019). Systems-level network modeling of Small Cell Lung Cancer subtypes identifies master regulators and destabilizers. PLoS Comput Biol 15, e1007343.

Wu, Q et al. (2021). YAP drives fate conversion and chemoresistance of small cell lung cancer. Sci Adv 7, eabg1850.

Yan, W, Chung, C-Y, Xie, T, Ozcek, M, Nichols, TC, Frey, J, Udyavar, AR, Sharma, S, and Paul, TA (2021). Intrinsic and acquired drug resistance to LSD1 inhibitors in small cell lung cancer occurs through a TEAD4-driven transcriptional state. Mol Oncol, in press.

Zhang, W et al. (2018). Small cell lung cancer tumors and preclinical models display heterogeneity of neuroendocrine phenotypes. Transl Lung Cancer Res 7, 32–49.

Zhou, JX, and Huang, S (2011). Understanding gene circuits at cell-fate branch points for rational cell reprogramming. Trends Genet 27, 55–62.

